# Class II UvrA protein Ecm16 requires ATPase activity to render resistance against echinomycin

**DOI:** 10.1101/2022.02.02.478902

**Authors:** Amanda Erlandson, Priyanka Gade, Chu-Young Kim, Paola Mera

## Abstract

Bacteria use various strategies to become antibiotic resistant. The molecular details of these strategies are not fully understood. We can increase our understanding by investigating the same strategies found in antibiotic-producing bacteria. In this work, we characterize the self-resistance protein Ecm16 encoded by echinomycin-producing bacteria. Ecm16 is a structural homolog of the Nucleotide Excision Repair (NER) protein UvrA. Expression of *ecm16* in the heterologous system *Escherichia coli* was sufficient to render resistance against echinomycin. Ecm16 preferentially binds double-stranded DNA over single-stranded DNA and is likely to primarily interact with the backbone of DNA using a nucleotide-independent binding mode. Ecm16’s binding affinity for DNA increased significantly when the DNA is intercalated with echinomycin. Ecm16 can repair echinomycin-induced DNA damage independently of NER. Like UvrA, Ecm16 has ATPase activity and this activity is essential for Ecm16’s ability to render echinomycin resistance. Notably, UvrA and Ecm16 were unable to complement each other’s function. Increasing the cellular levels of UvrA in *E. coli* was insufficient to render echinomycin resistance. Similarly, Ecm16 was unable to repair DNA damage that is specific to UvrA. Together, our findings identify new mechanistic details of how a refurbished DNA repair protein Ecm16 can specifically render resistance to the DNA intercalator echinomycin. Our results, together with past observations, suggest a model where Ecm16 recognizes double helix distortions caused by echinomycin and repairs the problem independently of NER.

## IV. Main Text

### (a) INTRODUCTION

The current crisis with antibiotic resistance has become one of the biggest public health challenges of our time (CDC). Bacteria can acquire antibiotic resistance through various mechanisms that include the upregulation of efflux pumps, modification of antibiotics, and modification and protection of the antibiotic target. Notably, many of these antibiotic resistance mechanisms have also been found in bacteria that are themselves producers of antibiotics, which are utilized for self-protection (1, 2). Some genes involved in these antibiotic resistance mechanisms have been proposed to originate from antibiotic-producing bacteria shared with pathogenic bacteria through transformation, transduction, or conjugation (3–6). This potential sharing of genetic information highlights the importance of identifying the mechanistic details of self-resistance found in antibiotic-producing bacteria.

About 80% of our known antibiotics are produced by *Streptomyces spp*. (7). One such antibiotic produced by *Streptomyces* is echinomycin, the first identified DNA bis-intercalator (8). Echinomycin, a member of the quinomycin family of antibiotics, contains two planar quinoxaline chromophores connected by a cyclic octadepsipeptide (9–11) (Figure 1A). Quinomycins have been shown to possess antimicrobial, antiviral, and antitumor activities (12–16). Echinomycin was originally discovered in *Streptomyces echinatus* but has since been found to be produced by several species of *Streptomyces* (17, 18). In one of these species, *Streptomyces lasolacidi* (formerly known as *S. lasaliensis)*, the echinomycin biosynthetic gene cluster lies within a 36 Kilobase region of a 520 Kilobase giant linear plasmid (19–21). This gene cluster consists of eight genes involved in quinoxaline-2-carboxylic acid synthesis, five genes for octadepsipeptide backbone synthesis (21), and five genes that encode proteins proposed to have regulatory or unknown functions (18). One of the genes of unverified function, *ecm16*, encodes a homolog of the prokaryotic nucleotide excision repair (NER) protein UvrA, which has been proposed to serve as a self-resistance protein against echinomycin (21). Interestingly, other DNA intercalator drugs like daunorubicin and triostin A also include a UvrA-homolog encoding gene in their biosynthetic clusters: *drrC* and *trsM* (1, 21–25). DrrC has been shown to render self-protection in *Streptomyces peucetius* against daunorubicin (26, 27). However, mechanistic understanding of how DrrC renders resistance against daunorubicin remains limited.

**Figure 1.**
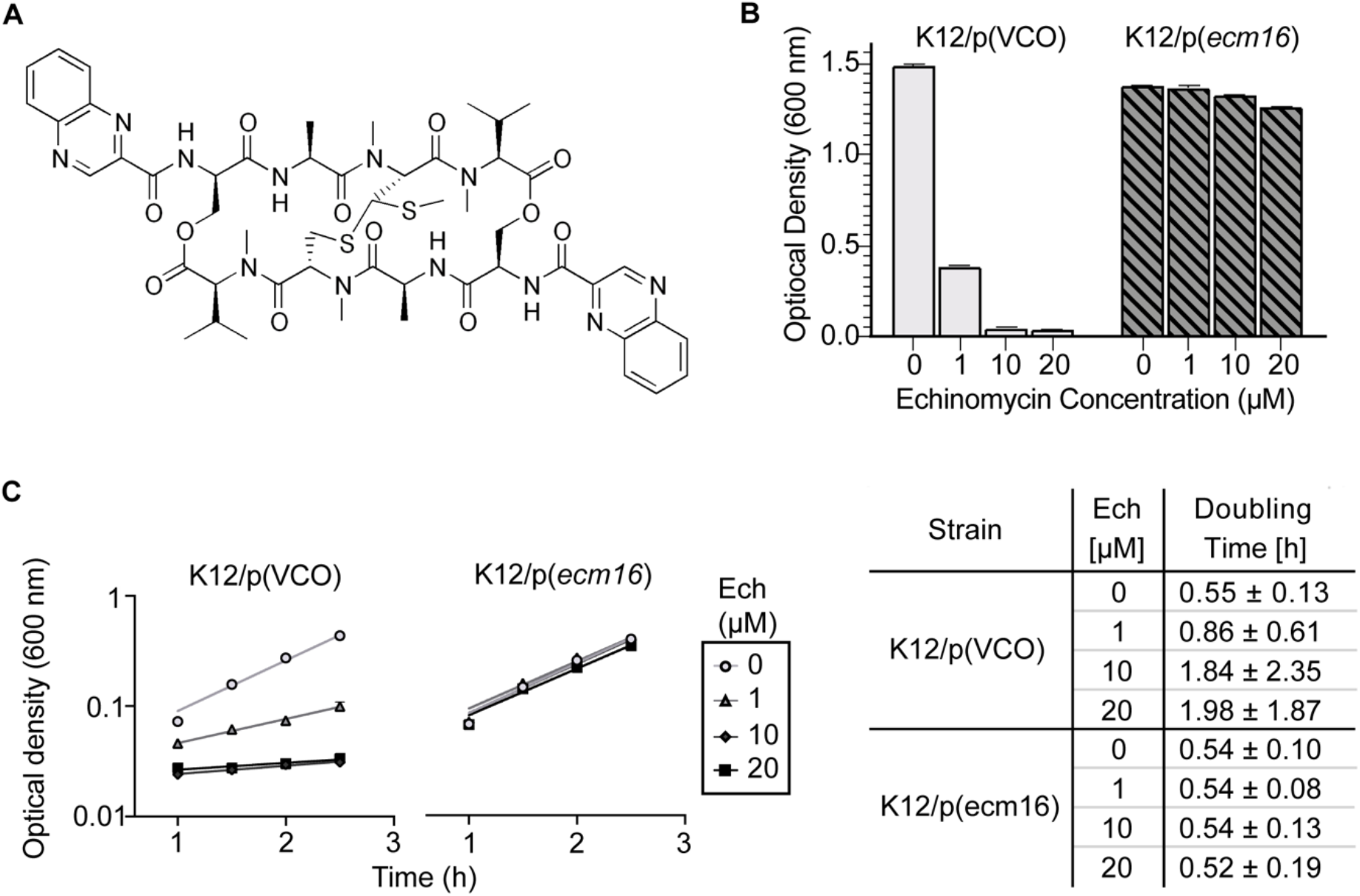
Expression of *ecm16* gives resistance against echinomycin. **(A)** Structure of echinomycin. **(B)** Optical density after 6 hours. Exposure to echinomycin results in reduction of *E. coli* growth in liquid media. Maximum optical densities were determined after 6 h exposure to echinomycin concentrations ranging from 1 μM to 20 μM. Echinomycin was added at time zero to cultures at 0.2 OD_600nm_. *E. coli* K12 strains with pBAD vector-control-only (VCO) or pBAD-*ecm16* (p(*ecm16*) were used for comparison. Error bars represent standard error for duplicate replicates of one trial, analysis is representative of three independent trials. **(C)** Exponential state growth curves of K12/pVCO and K12/p(*ecm16*) on a semi-logarithmic plot. Cultures were grown in rich media (LB) supplemented with echinomycin (0 to 20 μM) and arabinose inducer (0.2 %). Error bars represent standard error of the mean (SEM) of duplicate replicates. All results shown are representative of three independent replicates.

UvrA is part of the nucleotide excision repair (NER) system responsible for repairing diverse types of DNA damage in bacteria, including pyrimidine dimers (28, 29), unpaired T and G residues (30), backbone modifications such as single nucleotide gaps and nicks (26, 31–33), and damage conferred by anthramycin (31, 34, 35), cholesterol adducts (36), and fluorescein (33). The NER protein involved in recognizing DNA damage is UvrA, a dimeric protein with a total of four ATP-binding sites, two UvrB-binding domains, and a single DNA binding groove (37–41). ATPase activity conferred by the ATP-binding domains is necessary for UvrA’s interaction with DNA, dissociation of dimer into monomers, and interaction with UvrB (41, 42). Two models have been proposed regarding detection of damaged DNA: (1) UvrA scans DNA alone until it detects a lesion, at which point it stalls and complexes with UvrB (43) and (2) a UvrA2-UvrB2 complex searches and locates damaged DNA (44, 45). In either model, UvrA is the first protein of the NER system to bind DNA and the one responsible for promoting conformational changes of DNA and UvrB for both to bind to each other (46). Defining the molecular mechanism used by UvrA to recognize the wide range of DNA damage types has remained elusive in the field.

In addition to the canonical UvrA, several bacterial phyla (including *Actinobacteria, Firmicutes, Bacteroidetes* and *Proteobacteria)* also express a class II UvrA (47). The UvrA homologs found in *Streptomyces* that produce DNA intercalator drugs are class II UvrA (referred here as UvrA2) (1, 48). The specific functions of UvrA2 proteins in various bacterial species remain unclear. For instance, in *D. radiodurans*, expression of *uvrA* and *uvrA2* gene is upregulated following ionizing radiation (49). However, deletion of the *uvrA2* gene had no effect on the cell’s UV sensitivity (50). In *Xanthomonas axonopodis* and *Pseudomonas putida*, the double knockout Δ*uvrA* Δ*uvrA2* displayed slightly higher sensitivity to high UV radiation compared to the single knockout Δ*uvrA* suggesting that UvrA2 can contribute, although at a minor level, to the repair of UV-induced DNA damage (51). In *P. putida*, UvrA2 was proposed to also be involved in mutagenesis mechanisms during stationary growth phase (52). Overall, our knowledge about the specific roles that class II UvrA proteins play in bacteria and whether they can substitute the activity of the canonical UvrA remain limited.

In this study, we present the *in vivo* and *in vitro* characterization of Ecm16, a UvrA2 protein from *Streptomyces lasalocidi* (1, 48). We demonstrate *in vivo* that the expression of *ecm16* is sufficient to render the echinomycin-sensitive *E. coli* resistant to relatively high levels of echinomycin (20μM). *In vitro*, we showed that, unlike UvrA, Ecm16 preferentially binds echinomycin-containing double-stranded DNA. Similar to UvrA, Ecm16 can hydrolyze ATP suggesting that UvrA2 type proteins require ATPase activity for detecting/repairing DNA damage. Furthermore, Ecm16’s drug resistance activity does not require any of the other components of the classical NER pathway. Ecm16 was unable to complement an *E. coli uvrA* knockout strain recovering from UV-induced DNA damage. Collectively, our work provides new insights into how class II UvrA proteins catalyze drug resistance independently of the evolutionarily related protein UvrA.

### (b) RESULTS

#### Expression of *ecm16* provides resistance against echinomycin

Although previously proposed, there is no experimental data in the literature confirming that Ecm16 can render resistance against echinomycin (21). To provide this evidence and characterize Ecm16, we used *E. coli* (K12) cells as a heterologous host system to express *ecm16*. The natively echinomycin-sensitive *E. coli* cells were transformed with a low-copy vector encoding *ecm16* under the arabinose inducible promoter [p(*ecm16*)]. In the absence of echinomycin, *E. coli* cells with vector-control-only (VCO) reached saturation (OD_600nm_ ~1.0) within a 6 h growth period (Figure 1B). The same VCO cells grown in the presence of 1 μM echinomycin only reached ~0.35 OD_600nm_ within the same 6 h period. Almost no detectable growth was observed for VCO cells at echinomycin concentration of 10 μM or higher. In contrast, *E. coli* cells expressing *ecm16* reached equivalent maximal densities when grown in the presence or in the absence of supplemented echinomycin. Using growth curve analyses, we determined the effect of *ecm16* expression on doubling rates (Figure 1C). In the absence of echinomycin, *E. coli* with or without expression of *ecm16* were able to double every ~0.54 h under our growth conditions. However, in the presence of echinomycin, the doubling rate of VCO cells increased more than 7-fold. Remarkably, cells expressing *ecm16* were able to maintain their normal doubling rate up to levels of 20 μM echinomycin.

*E. coli* adopt filamentous shapes in response to a variety of stressful environments, including DNA damage and exposure to antibiotics (53–56). To determine whether Ecm16 can prevent cellular filamentation, we analyzed the cell length of *E. coli* after exposure to echinomycin. The average cell length of *E. coli* was determined to be *~*2.5 μm when grown in the absence of any drug. However, the average cell length increased six-fold (~14.9 μm) when grown in the presence of echinomycin (5 μM) (Figure 2). Exposure to echinomycin resulted in cells with broad cell length distribution and maximum lengths reaching up to ~54 μm. We observed that the expression of *ecm16* alone in the absence of any echinomycin, resulted in an average cell length that was longer (~3.8 μm) compared to the VCO (~2.5 μm). However, when echinomycin was supplemented to cells expressing *ecm16*, cell length was not altered compared to the no echinomycin control treatment. In summary, our *in vivo* analyses revealed that the expression of *ecm16* in *E. coli* cells results in protection against the toxicity caused by echinomycin exposure.

**Figure 2.**
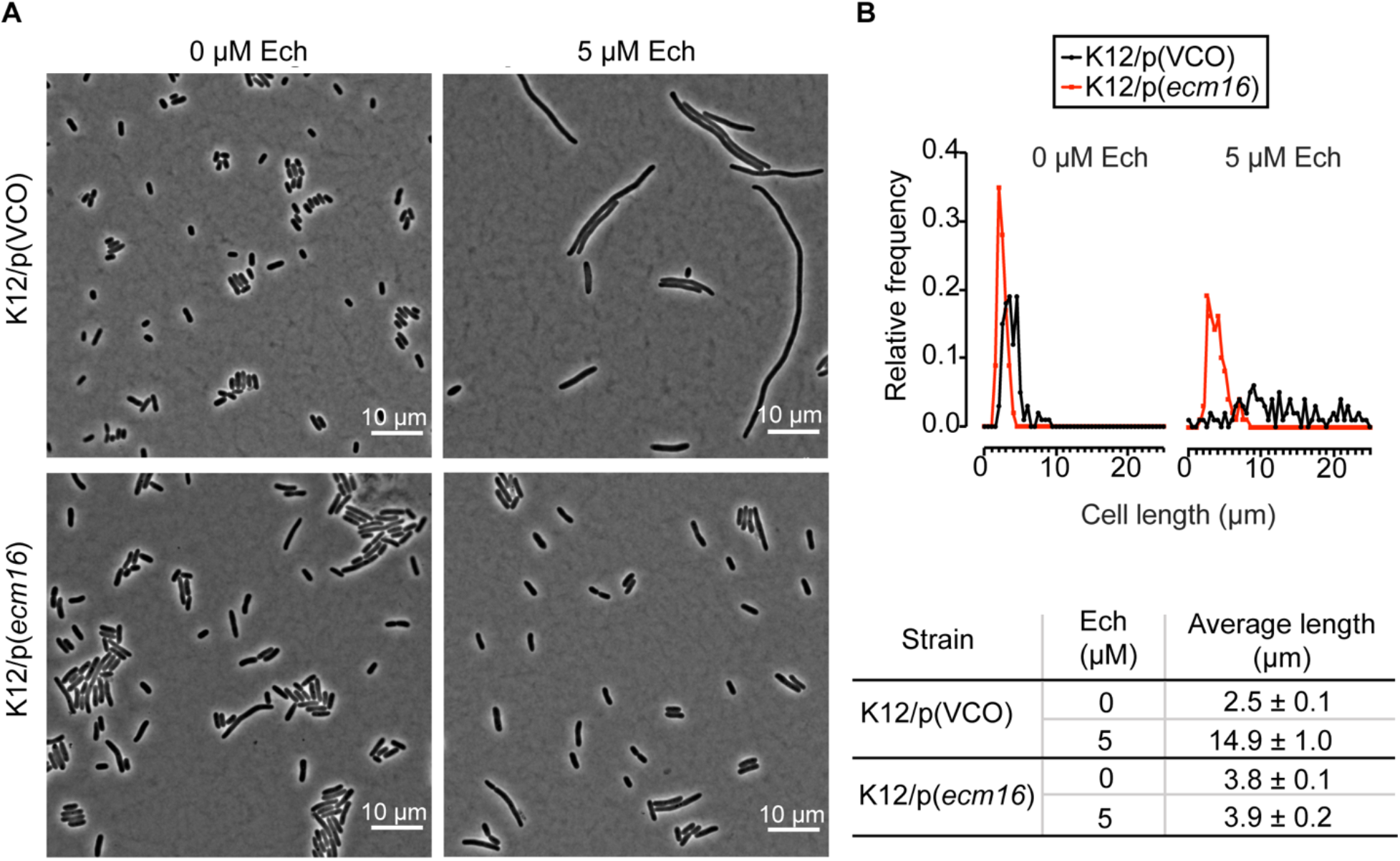
*E. coli* cells expressing *ecm16* do not filament after echinomycin treatment. **(A)** Phase contrast microscopy of strains K12/p(VCO) (top panels) and K12/p(*ecm16*) (bottom panels) grown in the presence of inducer (arabinose 0.2 %) supplemented with or without 5 μM echinomycin (ech). Cells were grown in the presence of echinomycin for 5 h and subsequently spotted on 1 % agarose minimal media pads for imaging. Scale bar = 10 μm. **(B)** Cell size distribution and average cell length and standard error for 100 cells per condition are shown. 7% of cells displayed cell lengths over 25 μm (not included in plot). Cell lengths were measured using the software MicrobeJ. Representatives of three independent replicates are plotted. Cell lengths were binned in 0.5 μm increments and plotted using Prism.

#### Ecm16 binds preferentially to double-stranded DNA-echinomycin

We next investigated how the local DNA structure influences Ecm16’s ability to recognize its substrate. The NER UvrA from *E. coli* and *Mycobacterium tuberculosis* has been shown to exhibit stronger affinity for single-stranded DNA (ssDNA) versus double-stranded DNA (dsDNA) (57, 58). Using electrophoretic mobility shift assay, we characterized the DNA binding activity of purified recombinant Ecm16 to various DNA substrates (Figure 3A). Our data revealed that unlike UvrA, Ecm16 displays stronger binding affinity to dsDNA compared to ssDNA suggesting a potential mechanistic difference in damage recognition and/or repair (Figure 3B). The substrate affinity of Ecm16 was further increased when dsDNA was combined with echinomycin (Figure 3AB). Given that echinomycin intercalates DNA with a preference for CpG sites (10, 59), we tested whether Ecm16 displays stronger affinity for specific DNA sequences. To determine whether Ecm16 displayed any DNA preference, we analyzed different combinations of DNA sequences using electrophoretic mobility shift assays where the levels of Ecm16 were titrated. We varied the percent of GC and AT content of the DNA substrate (Figure 3C). Our results revealed no significant differences in Ecm16’s substrate binding with varying GC/AT content, suggesting that Ecm16 primarily interacts with the backbone, and not the bases, of DNA. These data indicate that Ecm16 has substrate specificity and/or higher binding affinity for dsDNA-echinomycin and binds DNA in a nucleotide-independent mode. Collectively, these data suggest that Ecm16 does not explicitly bind regions prone to echinomycin binding but instead Ecm16 may scan dsDNA to detect lesions caused by echinomycin.

**Figure 3.**
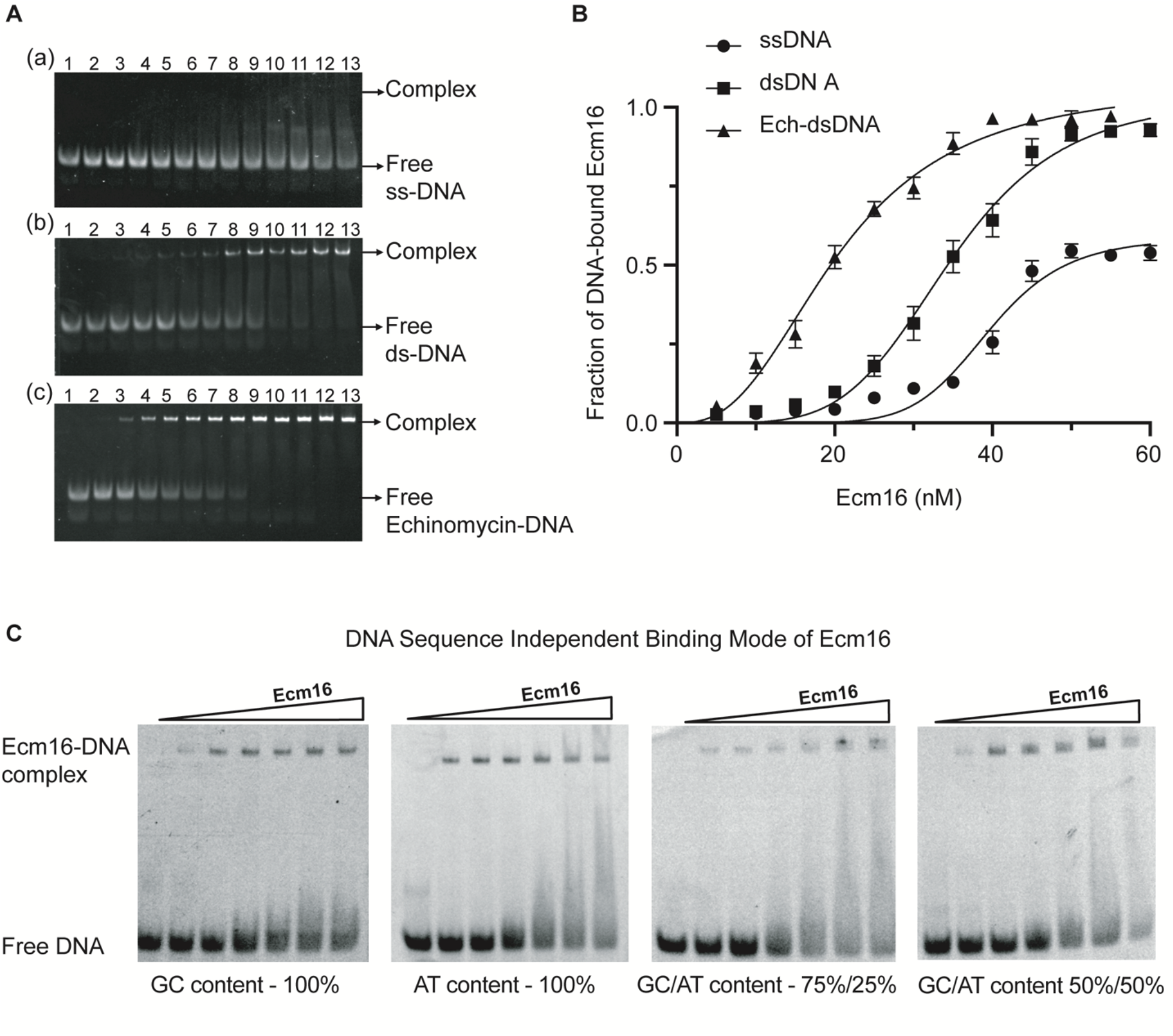
DNA binding activity of Ecm16. **(A)** Reaction mixtures contained 0.25 nM DNA substrate in the absence (lane 1) or presence of 5 to 60 nM Ecm16 with increment of 5 nM (lanes 2-13), (a) ssDNA; (b) dsDNA; (c) Echinomycin-DNA. **(B)** Fraction of ssDNA (panel (a)), dsDNA (panel (b)), Echinomycin-DNA (panel (c)) bound to Ecm16 is plotted against the indicated amounts of Ecm16. ssDNA (filled circles); dsDNA (filled squares); Echinomycin-DNA (filled triangles). Each point on the curves represents the mean of three separate experiments. **(C)** DNA binding activity of Ecm16 in presence of 100% GC and AT, 75%/25% and 50%/50% GC/AT composition 0.25 nM DNA substrates in the absence (lane 1) or presence of 5 to 30 nM Ecm16 with increment of 5 nM (lanes 2-7).

#### Ecm16 displays ATPase activity that is essential for echinomycin resistance

The ATPase activity of UvrA is required for UvrA’s ability to recognize DNA damage and initiate the repair mechanism (42, 60). Ecm16, like other UvrA homolog proteins, has high conservation of the amino acids involved in ATP binding/hydrolysis, including those found in Walker A, Walker B, and the alpha-helical ABC signature sequence (Supplemental Figure 1). To determine whether Ecm16 displays ATPase activity and whether this activity is involved in protection against echinomycin, we first characterized Ecm16’s ability to hydrolyze ATP *in vitro*. To do that, we used a coupled assay in which the substrate, 2-amino-6-mercapto-7-methylpurine riboside, is enzymatically converted to ribose 1-phosphate and 2-amino-6-mercapto-7-methylpurine by purine nucleoside phosphorylase in the presence of inorganic phosphate (Figure 4A). Ecm16 shows basal level ATP hydrolysis activity in the absence of any substrate, a characteristic which is also observed in UvrA (57). The addition of dsDNA increased Ecm16’s ATPase specific activity ~10-fold, while addition of echinomycin-dsDNA complex increased the ATPase activity ~200-fold (Figure 4). We next determined whether the observed ATPase activity of Ecm16 was involved in Ecm16’s ability to render echinomycin resistance *in vivo*. To do that, we targeted a conserved residue (Lys526) in UvrA-type proteins that has been shown to be essential for discriminating between native and damaged DNA (61–64). To determine whether Ecm16 requires its ATPase activity *in vivo*, we engineered an Ecm16 variant with the conserved residue Lys526 changed to an alanine and analyzed the ability of an *E. coli* strain expressing this variant (Ecm16^K526A^) to confer echinomycin resistance. Our data revealed that *E. coli* cells exclusively expressing the variant Ecm16^K526A^ failed to rescue cells from echinomycin toxicity (Figure 4B). These data suggest that Ecm16 may use ATP hydrolysis for scanning DNA for damage recognition and/or repair.

**Figure 4.**
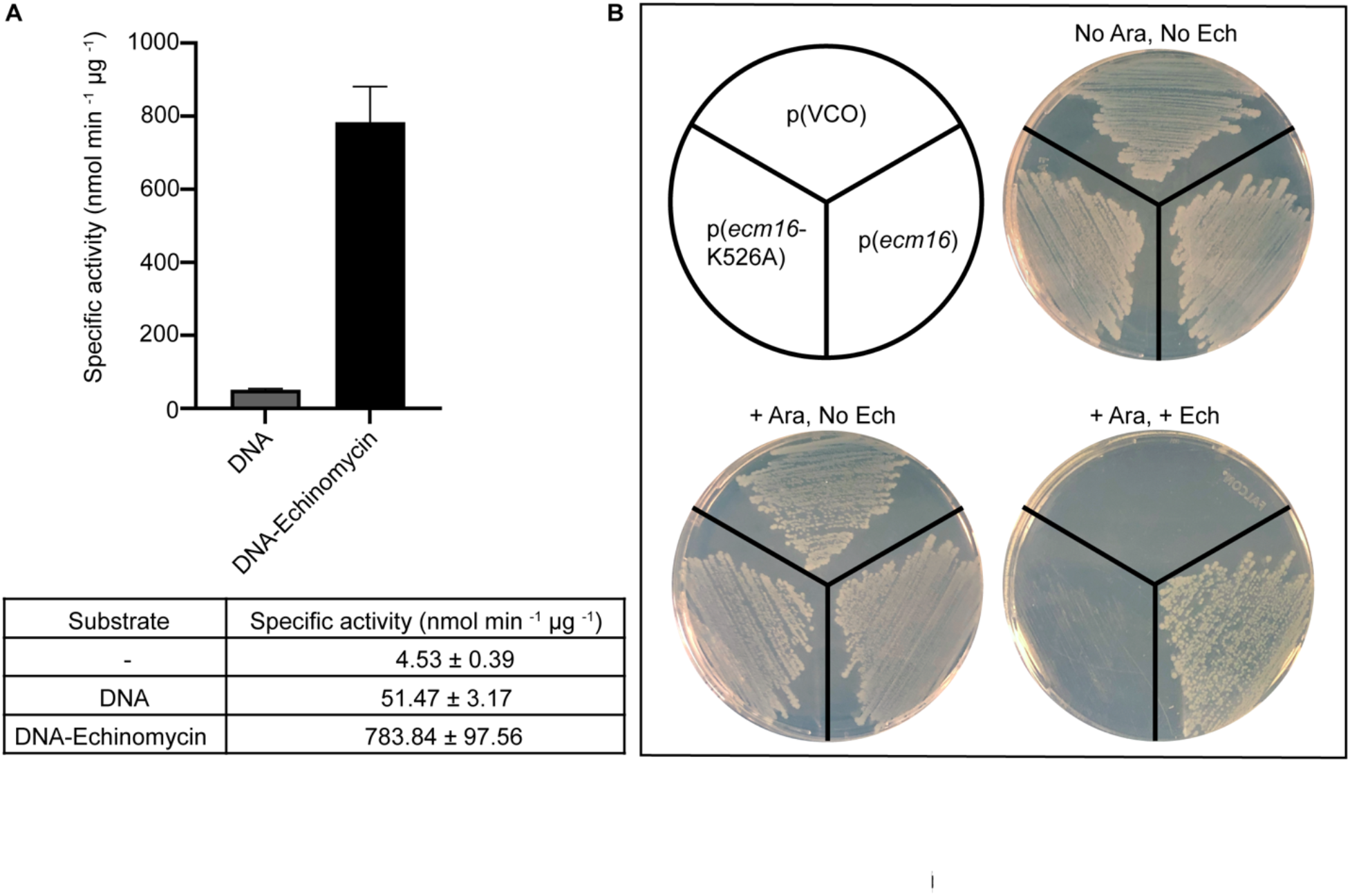
**(A)** Specific activity of 0.2 μM purified Ecm16 in the presence or absence of 1 μM DNA and DNA-echinomycin substrates. Error bars represent standard error of three independent experiments. The reaction mixture contained 2-amino-6-mercapto-7-methylpurine (MESG) and purine nucleoside phosphorylase, 1 mM ATP and 10 mM MgCl_2_. A phosphate standard was measured to calibrate the UV absorbance signal to the amount of inorganic phosphate release. **(B)** Streak plates containing *E. coli* cultures containing a plasmid with WT *ecm16* (p(*emc16*)), *ecm16* ATP-binding variant (p(*emc16-K526A*)), or vector control (p(VCO)). Cultures were grown overnight and normalized to 0.5 OD_600_ before streaking on LB ampicillin plates containing 0.2% arabinose or 1 μM echinomycin (with 0.2% arabinose) and were incubated at 37°C overnight. Plates are representative of 3 independent trials.

#### Ecm16’s echinomycin resistance activity does not require components of the NER system

Based on similarities between Ecm16 and UvrA, we examined whether Ecm16’s mechanism of protective action against echinomycin resembles the activity of the DNA repair protein UvrA. In *E. coli*, UvrA or UvrA2-UvrB2 detects DNA damage and recruits the rest of the NER system (UvrB/C/D) (65–68). To address the possibility that Ecm16’s ability to render cells echinomycin resistance requires the function of NER components, we analyzed *E. coli* strains with knockouts of each individual component (*uvrA, uvrB, uvrC*, and *uvrD*; Keio Collection) (69) (Figure 5A). To each of these strains, we transformed the replicating plasmid encoding *ecm16* or VCO. We tested the ability of these strains to grow in the presence of echinomycin concentrations that were tolerated by cells expressing *ecm16*. Our growth analyses revealed that cells encoding the NER system and cells without components of the NER system displayed similar doubling rates when grown in the presence of echinomycin. Ecm16 was able to provide the same level of echinomycin resistance in the absence of NER.

**Figure 5.**
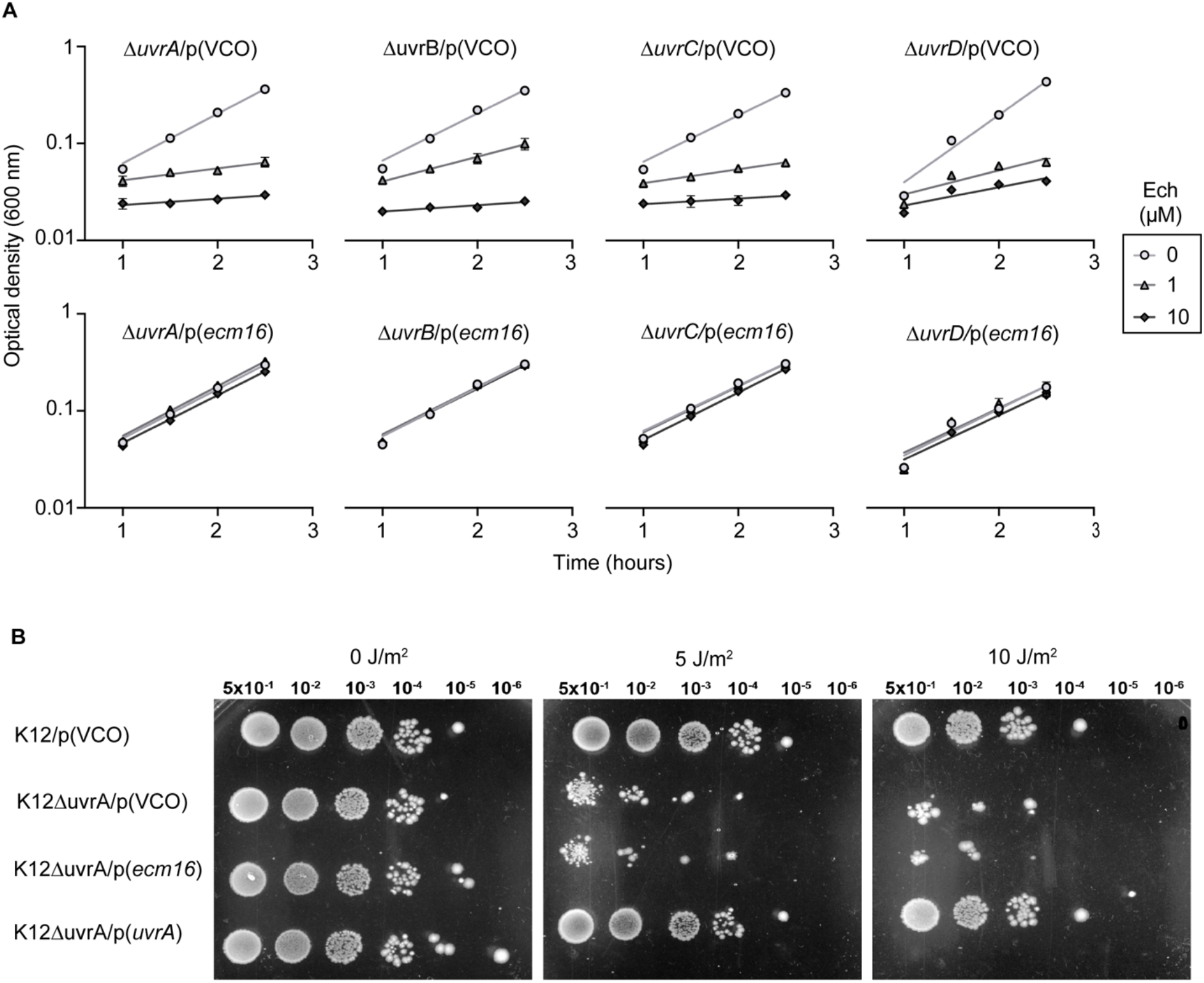
Ecm16 is not associated with classic NER function. **(A)** Exponential state growth curves (n = 3) of *E. coli* K12 strains with different components of the native nucleotide excision repair system (NER) knocked out (Δ*uvrA*, Δ*uvrB*, Δ*uvrC* or Δ*uvrD*) and carrying vector-control-only or vector encoding *ecm16*. Cultures (2 mL) were set to OD_600nm_ ~ 0.2 in rich media (LB) supplemented with the inducer (0.2% arabinose) and varying concentrations of echinomycin (0 - 10 μM). Growth was monitored by measuring the absorbance at 600 nm every 30 minutes. *E. coli* K12 NER knockouts show similar patterns of resistance to echinomycin in the presence of inducer for the expression of ecm16. Results are shown for exponential growth phase with exponential trend line, error bars represent SEM of duplicate replicates. All results shown are representative of three independent replicates. **(B)** Colony Forming Unit (CFU) assays after UV radiation. 5 μL of cultures grown to OD_600nm_ ~0.2 were serial diluted (dilution factor = 5 x 10^-1^ to 10^-6^) and spotted on LB plates supplemented with 0.2% arabinose. Cultures included the following strains: control sample (top row) are wild-type cells with vector-control-only, test samples (middle 2^nd^ and 3^rd^ rows) are strains with the native *E. coli’s* uvrA gene knocked out and with vector-control-only or vector encoding *ecm16*, complementation control strain (bottom row) encodes *E. coli’s* native *uvrA* gene in the same pBAD vector. Freshly spotted plates were exposed to UV radiation and then incubated at 37 °C for 18 h prior to imaging. The data shown are a representative of three independent replicates.

#### Ecm16 and UvrA cannot complement each other’s function

In *E. coli*, the activity of UvrA is essential for repairing various types of DNA damage including thymine dimers resulting from exposure to UV radiation (70). To determine whether Ecm16 can complement UvrA’s activity, we examined the ability of various strains to recover from UV radiation by performing Colony Forming Unit (CFU) assays (Figure 5B). *E. coli* cells with their native *uvrA* gene and VCO can effectively recover from 10 J/m^2^ of UV radiation exposure as evidenced by the similar number of CFUs compared to the no-radiation control. However, the cell’s ability to recover from UV radiation is significantly reduced when *E. coli’s* native *uvrA* gene is knocked out. Once *uvrA* was expressed *in trans* from a replicating plasmid under the control of the arabinose inducible promoter, the ability to repair DNA damage was fully recovered. These data are in accordance with previous analyses of *uvrA* knockout strains (71). However, when the gene encoding for Ecm16 was expressed *in trans* from the same replicating plasmid controlled by the arabinose promoter, *E. coli* cells were unable to recover from UV radiation. The number of CFUs of the strain expressing *ecm16* was as diminished as the CFUs observed on the *uvrA* knock out strain. The same trend of UV resistance was also observed among the *E. coli* strains when lower levels of radiation (5 J/m^2^) were used. The inability of Ecm16 to complement a *uvrA* knockout suggests that Ecm16 is unable to recognize thymine dimers and/or recruit the rest of the NER system for proper DNA repair.

Given that *E. coli* K12 cells display sensitivity to echinomycin suggested that UvrA is unable to protect from echinomycin toxicity. To determine whether increasing the cellular levels of UvrA would result in protection against echinomycin, we constructed an *E. coli* strain with two copies of *uvrA:* its native copy and a second copy expressed *in trans* from a replicating plasmid (same vector used for *ecm16* expression). Cells expressing *uvrA* from the inducible promoter grown in the presence of echinomycin display the same growth defect as the empty vector control (Supplemental Figure 2). These results confirm that UvrA and Ecm16 cannot complement each other’s function due to either differences in damage detection and/or repair.

### (c) DISCUSSION

In this study, we provide *in vivo* and *in vitro* characterization of Ecm16, a class II UvrA protein. Our data demonstrate that expression of *ecm16* is sufficient to confer echinomycin resistance to an otherwise echinomycin sensitive bacterium, *E. coli* K12. We show that ATP hydrolysis of Ecm16 is required for its ability to render resistance. Furthermore, Ecm16 was able to confer echinomycin resistance to host cells which are deficient in UvrA, UvrB, UvrC, or UvrD, indicating that Ecm16 does not depend on the NER machinery for its function. Finally, our data reveal that despite the sequence similarity between UvrA and Ecm16, they are unable to complement each other’s function. Collectively, our data expand our current understanding of the various mechanism that UvrA2 proteins utilize for their function.

Our work provides further insights into the potential mechanism used by Ecm16 to render resistance from echinomycin toxicity. The fact that Ecm16 binds DNA-echinomycin specifically implies that Ecm16’s mode of action relies on Ecm16 binding DNA, and not detoxifying the cell by sequestering echinomycin away from DNA. Furthermore, UvrA recognizes DNA lesions and promotes the recruitment of the rest of the NER system to excise the damaged DNA. However, Ecm16 retained its ability to render echinomycin resistance in the absence of any of the NER proteins. This observation posits two potential models where Ecm16 can either work with other housekeeping proteins found in *E. coli* or Ecm16 can work alone. The former model is plausible given that UvrA has been shown to work collaboratively with non-NER proteins (72, 73). In the scenario where the latter model is correct, we envision Ecm16 to use ATP hydrolysis to unwind, stretch, or compress DNA thereby dislodging echinomycin from DNA. UvrA has been proposed to subject DNA to a ‘stress test’ where UvrA unwinds and stretches-compresses DNA during the detection of damage (74). Ecm16 could be using a version of this ‘stress test’ mechanism but in doing that it would potentially release the echinomycin bound and thus repair the damage of DNA. We are currently testing between these two potential models.

Although multiple models have been proposed to explain how the NER system recognizes DNA damage, conclusively addressing the mechanism of recognition remains a major gap in our knowledge in both bacterial and eukaryotic NER systems. We propose that the understanding of the mechanism of substrate recognition by Ecm16 can provide insights into how the NER system recognizes its substrates. This is particularly the case given that UvrA and Ecm16 were unable to promote each other’s function. Ecm16 and UvrA may be able to detect different types of distortions of the DNA double helix. UvrA has been proposed to distinguish between native and damaged DNA by sensing changes in various properties of DNA (structural, electrostatic, dynamics, stability, flexibility) (46). Our data revealed that UvrA is unable to repair echinomycin-induced damage possibly because the distortions of DNA caused by echinomycin are not recognized by UvrA or the NER system is unable to repair the damage. Figuring out details of either mechanism would advance our understanding of NER and DNA damage recognition and repair. Furthermore, UvrA and Ecm16 seem to also have different preferences between ssDNA and dsDNA that can potentially illustrate differences in damage recognition. Further work on Ecm16 will advance not only our understanding of how antibiotic-producer bacteria become antibiotic resistant but also advance understanding of the conserved NER system.

### (d) EXPERIMENTAL PROCEDURES

#### Bacterial growth conditions

*E. coli* strains were grown in Luria Bertani broth (LB) (Fisher BioReagents, NJ, U.S.A.). All cultures grown in liquid media were grown at 37°C with orbital shaking at 200 rpm. Strains cultured on solid media were plated on LB agar (Fisher BioReagents, NJ, U.S.A.) and incubated at 37°C. Ampicillin (amp) resistant strains were grown with 50 μg amp/mL liquid media (Ampicillin sodium salt prepared in H_2_O, Amresco, Ohio, U.S.A.) and 100 μg amp/mL solid media. Kanamycin (kan) resistant strains were treated with 30 μg kan/mL for liquid media (kanamycin sulfate prepared in H_2_O, IBI Scientific, NJ. U.S.A.), and 50 μg kan/mL for solid media. For induction, 0.2% L-(+)-Arabinose (Alfa Aesar, G.B.) was added to either liquid or solid media. Strains grown with echinomycin were treated with 1 μM-20 μM of echinomycin diluted in H_2_O from a 908 μM echinomycin (Sigma, U.S.A.) solution prepared in methanol (Fisher Chemical, NJ, U.S.A.). All strains and primers used for this study are listed in Supplemental Tables S1 and S2.

#### Growth analyses

Cultures were grown in liquid media with ampicillin overnight from frozen stocks, incubated at 37°C and 200 rpm. Saturated cultures were diluted with fresh medium and induced with a 0.2% arabinose solution for 60 minutes and used to inoculate 2 mL duplicate replicate samples at a starting optical density of 0.02 OD_600_ in 13 mm glass tubes. Cultures were grown in rich liquid media (LB), with Ampicillin and 0.2% Arabinose, incubated at 37°C and 200 rpm. Optical density readings were taken every 30 minutes for 6 hours using Thermo UV-Vis Spectrophotometer. Maximum growth in echinomycin was determined by evaluating serial dilutions of echinomycin in H_2_O when added to LB media inoculated with K12 *E. coli* culture, using 1 μM to 20 μM ech. K12/pBAD (vector control) and K12/pBAD-*ecm16* were grown overnight from frozen stocks at 37°C and 200 rpm, and prepared as in growth curves. Endpoint readings were taken after 6 hours of incubation.

#### Imaging

K12 strains were grown from frozen stocks and used to inoculate 3 mL of LB/Amp/0.2% arabinose to 0.2 OD_600_. Cells were induced with 0.2% arabinose 30 minutes prior to dilution. Following inoculation, cultures were treated with 5μM echinomycin and grown for 5 hours. Cultures were spotted on 0.2% agarose (Ultrapure, Invitrogen, Spain) discs, and visualized with phase contrast microscopy using Zeiss Axio Observer 2.1 inverted microscope with a Plan-Apochromat 100×/1.40 Oil Ph3 M27 (WD=0.17 mm) objective, AxioCam 506 mono camera and ZEN software. Cell size was calculated using ImageJ/FIJI with MicrobeJ (75–78)

#### Colony Forming Units

Cultures were grown overnight incubated at 37°C and 200 rpm in liquid media with ampicillin and used to inoculate samples to 0.3 OD_600_ the following day. 0.2% arabinose was added to each sample and cells were incubated for 2 hours. Cells were then serially diluted by factors of 10 and 5μL of cultures were spotted on LB/Ampicillin plates with 0.2% Arabinose. Plates were exposed to ultraviolet radiation (nm) at 5 J/m^2^ or 10 J/m^2^ or no UV as a negative control.

#### Purification of Ecm16

Recombinant proteins were expressed by transforming *E. coli* BL21 (de3) (Novagen, Merck Millipore) host strains with the recombinant pET28a constructs encoding Ecm16 wildtype. Cells were cultured at 37°C in 5 ml of LB medium supplemented with kanamycin to an optical density of ~0.6-0.8 at 600 nm. Protein expression was performed by inducing the cells with 0.1 mM isopropyl-β-D-thiogalactoside, and growth was continued for a further 16 h at 18°C. Cell pellet was resuspended in lysis buffer containing 50 mM HEPES pH 7.5, 200 mM NaCl, 10% glycerol, 10 mM Imidazole, 1 mM Phenylmethylsulphonyl fluoride (PMSF), 10 μg/ml DNase and 10 mM MgCl_2_. Cells were disrupted using using sonication (Branson Ultrasonics) and cell debris was discarded by centrifugation at 15,000 *g* for 45 minutes at 4 °C. Cleared cell lysate was applied on 5 ml His-Trap Crude column (GE Healthcare) equilibrated with loading buffer (50 mM HEPES, pH, 7.5 and 200 mM NaCl). The protein was eluted with a step gradient of imidazole (250 mM) using elution buffer (50 mM HEPES pH 7.5, 200 mM NaCl, 500 mM Imidazole). Ecm16 eluted from the nickel column was diluted up to 10-fold using dilution buffer (50 mM HEPES pH 7.5, 50 mM NaCl). The diluted protein was applied to 5 ml Hi-Trap Q HP anion-exchange column (GE Healthcare). Protein was eluted with 5 column volumes (CV) linear NaCl gradient from 50 mM to 500 mM concentration. The protein was concentrated up to 4-6 mg ml^-1^ and gel filtration was performed on Superdex 200 10/300 GL (GE Healthcare) equilibrated with storage buffer (50 mM HEPES pH 7.5, 50 mM NaCl). The eluted protein sample was concentrated up to 8-10 mg ml^−1^, shock-frozen in liquid nitrogen and stored at - 80°C.

#### Electrophoretic mobility shift assay

Stock solutions of echinomycin (908 μM) (Cayman Chemical) were prepared by dissolving the echinomycin powder in 100% methanol. DNA-echinomycin complex was allowed to form by incubating 1: 1.1 molar ratio of DNA:echinomycin for 15 min at room temperature. DNA-echinomycin was incubated at room temperature for 20 min with Ecm16 protein at concentrations ranging from 100 to 300 nM in 10 μl reactions containing binding buffer (50 mM HEPES pH 7.5, 50 mM NaCl). The reaction was terminated by adding 4% sucrose solution. The protein-DNA complex was electrophoretically separated at 4°C on 6% native polyacrylamide TBE gel (Invitrogen) at 75 V/cm. The gel was stained for 10 min in 1x TBE buffer containing 5 μL of SYBR gold nucleic acid gel stain (Invitrogen). The band intensity corresponding to the free DNA and protein-DNA were visualized by ultraviolet transillumination complex and the bands were quantified using ImageJ software (79).

#### ATPase activity assay

Adenosine triphosphate (ATP) was dissolved in storage buffer (50 mM HEPES pH 7.5, 50 mM NaCl), to obtain 100 mM stock concentration. Hydrolysis rate of ATP by Ecm16 was measured using a coupled enzyme assay where the substrate 2-amino-6-mercapto-7-methylpurine riboside (MESG) is enzymatically converted to ribose 1-phosphate and 2-amino-6-mercapto-7-methylpurine by purine nucleoside phosphorylase (PNP) in presence of inorganic phosphate (Pi). The reaction mixture (100 μL) contained 50 mM HEPES pH 7.5, 50 mM NaCl, 0.5 mg/ml BSA, 10 % glycerol, 10 mM MgCl_2_, 1 mM ATP, 0.2 mM MESG, 1x reaction buffer (50 mM Tris HCl pH 7.5, 10 mM MgCl2, 0.1 mM sodium azide), 1 U/ml PNP and 0.5 μM Ecm16 protein. The Reaction was carried by monitoring the increase in absorbance at 360 nm over a 60-minute period. The rate of ATP hydrolysis was calculated from the linear change in A360nm, with correction for MESG conversion to 2-amino-6-mercapto-7-methylpurine in absence of Ecm16 protein. The effect of DNA on ATP hydrolysis was studied by adding 2.5 μM 32-mer double or single-stranded DNA (containing single echinomycin binding site), 2.5 μM echinomycin antibiotic to the assay mixture.

## V. Acknowledgements

This work was supported by NIH RISE R25GM061222 (A.E.) and the University of Texas System STARs Award (C.-Y.K.).

## VI. Author Contributions

P.M. and C.-Y.K. conceived and supervised the project. P.G. conducted ATP hydrolysis and EMSA measurements. A.E. performed bacterial growth and microscopy studies. P.G., A.E., P.M. and C.-Y.K. wrote the manuscript.

