## Supplemental for "Class II UvrA protein Ecm16 requires ATPase activity to render resistance against echinomycin"

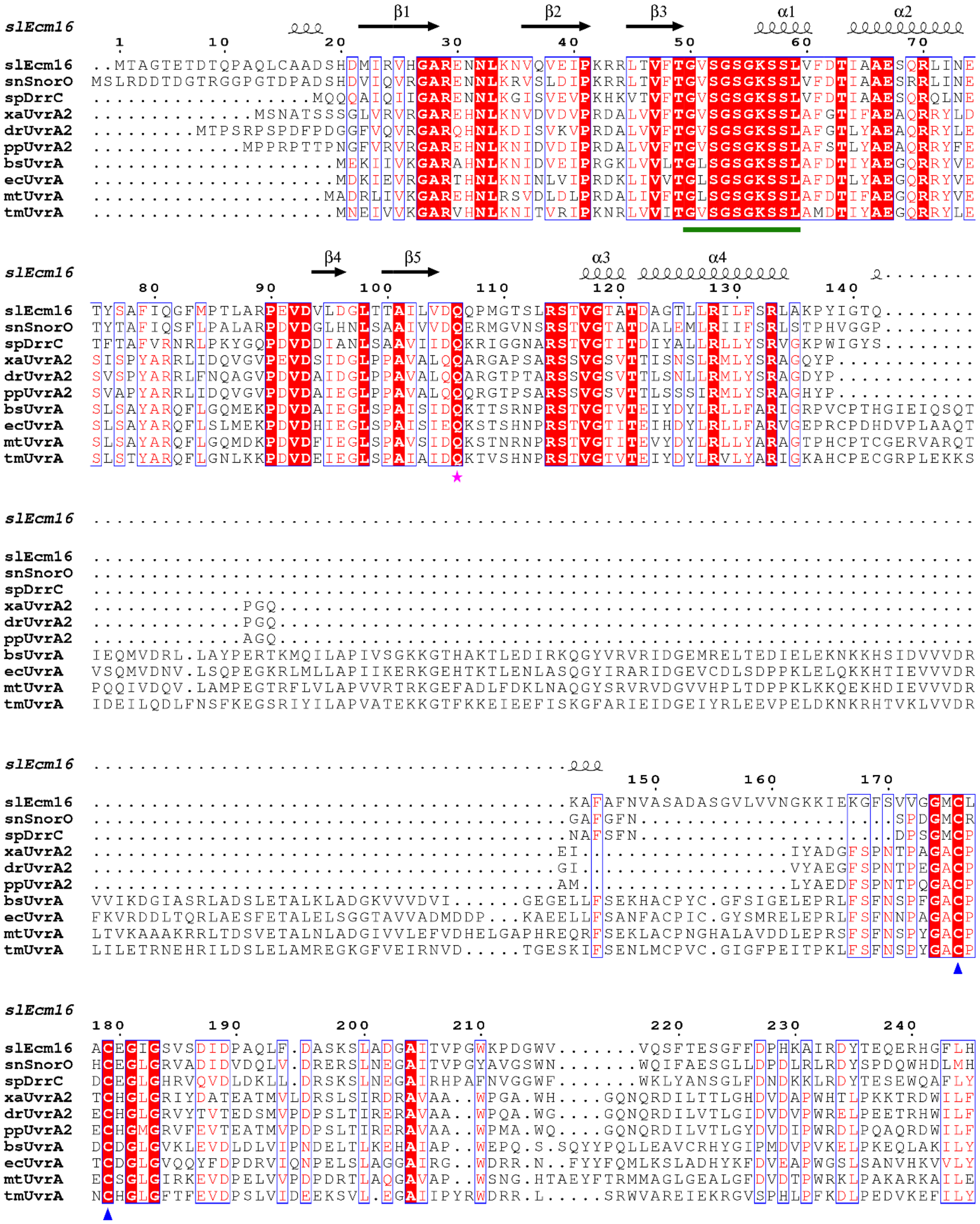
slEcm16
β1 β2 β3 α1 α2
1 10 20 30 40 50 60 70
slEcm16 ..MTAGTETDTQPAQLCAADSHDMIRVHGARENNLKNVQVEIPKRRLTVFTGVSGSGKSSLVFDTIAAESQRLINE
snSnorO MSLRDDTDGTRGGPGTDPADSHDVIQVRGARENNLKRVSLDIPKRRLTVFTGVSGSGKSSLVFDTIAAESRRLINE
spDrrC ...................MQQQAIQIIGARENNLKGISVEVPKHKVTVFTGVSGSGKSSLVFDTIAAESQRQLNE
xaUvrA2 ..............MSNATSSSGLVRVRGAREHNLKNVDVDVPRDALVVFTGVSGSGKSSLAFGTIFAEAQRRYLD
drUvrA2 .........MTPSRPSPDFPDGGFVQVRGARQHNLKDISVKVPRDALVVFTGVSGSGKSSLAFGTLYAEAQRRYLE
ppUvrA2 .............MPPRPTTPNGFVRVRGAREHNLKNIDVDIPRDALVVFTGVSGSGKSSLAFSTLYAEAQRRYFE
bsUvrA .....................MEKIIVKGARAHNLKNIDVEIPRGKLVVLTGLSGSGKSSLAFDTIYAEGQRRYVE
ecUvrA .....................MDKIEVRGARTHNLKNINLVIPRDKLIVVTGLSGSGKSSLAFDTLYAEGQRRYVE
mtUvrA ....................MADRLIVKGAREHNLRSVDLDLPRDALIVFTGLSGSGKSSLAFDTIFAEGQRRYVE
tmUvrA .....................MNEIVVKGARVHNLKNITVRIPKNRLVVITGVSGSGKSSLAMDTIYAEGQRRYLE
slEcm16
β4 β5 α3 α4
80 90 100 110 120 130 140
slEcm16 TYSAFIQGFMPTLARPEVDVLDGLTTAILVDQQPMGTSLRSTVGTATDAGTLLRILFSRLAKPYIGTQ........
snSnorO TYTAFIQSFLPALARPDVDGLHNLSAAIVVDQERMGVNSRSTVGTATDALEMLRIIFSRLSTPHVGGP........
spDrrC TFTAFVRNRLPKYGQPDVDDIANLSAAVIIDQKRIGGNARSTVGTITDIYALLRLLYSRVGKPWIGYS........
xaUvrA2 SISPYARRLIDQVGVPEVDSIDGLPPAVALQQARGAPSARSSVGSVTTISNSLRMLYSRAGQYP............
drUvrA2 SVSPYARRLFNQAGVPDVDAIDGLPPAVALQQARGTPTARSSVGSVTTLSNLLRMLYSRAGDYP............
ppUvrA2 SVAPYARRLIDQVGVPDVDAIEGLPPAVALQQQRGTPSARSSVGSVTTLSSSIRMLYSRAGHYP............
bsUvrA SLSAYARQFLGQMEKPDVDAIEGLSPAISIDQKTTSRNPRSTVGTVTEIYDYLRLLFARIGRPVCPTHGIEIQSQT
ecUvrA SLSAYARQFLSLMEKPDVDHIEGLSPAISIEQKSTSHNPRSTVGTITEIHDYLRLLFARVGEPRCPDHDVPLAAQT
mtUvrA SLSAYARQFLGQMDKPDVDFIEGLSPAVSIDQKSTNRNPRSTVGTITEVYDYLRLLYARAGTPHCPTCGERVARQT
tmUvrA SLSTYARQFLGNLKKPDVDEIEGLSPAIAIDQKTVSHNPRSTVGTVTEIYDYLRVLYARIGKAHCPECGRPLEKKS
slEcm16 ............................................................................
slEcm16 ............................................................................
snSnorO ............................................................................
spDrrC ............................................................................
xaUvrA2 .............PGQ............................................................
drUvrA2 .............PGQ............................................................
ppUvrA2 .............AGQ............................................................
bsUvrA IEQMVDRL.LAYPERTKMQILAPIVSGKKGTHAKTLEDIRKQGYVRVRIDGEMRELTEDIELEKNKKHSIDVVVDR
ecUvrA VSQMVDNV.LSQPEGKRLMLLAPIIKERKGEHTKTLENLASQGYIRARIDGEVCDLSDPPKLELQKKHTIEVVVDR
mtUvrA PQQIVDQV.LAMPEGTRFLVLAPVVRTRKGEFADLFDKLNAQGYSRVRVDGVVHPLTDPPKLKKQEKHDIEVVVDR
tmUvrA IDEILQDLFNSFKEGSRIYILAPVATEKKGTFKKEIEEFISKGFARIEIDGEIYRLEEVPELDKNKRHTVKLVVDR
slEcm16 .........................................
150 160 170
slEcm16 .........................................KAFAFNVASADASGVLVVNGKKIEKGFSVVGGMCL
snSnorO .........................................GAFGFN....................SPDGMCR
spDrrC .........................................NAFSFN....................DPSGMCP
xaUvrA2 ........................................EI..................IYADGFSPNTPAGACP
drUvrA2 ........................................GI..................VYAEGFSPNTPEGACP
ppUvrA2 ........................................AM..................LYAEDFSPNTPQGACP
bsUvrA VVIKDGIASRLADSLETALKLADGKVVVDVI.....GEGELLFSEKHACPYC.GFSIGELEPRLFSFNSPFGACP
ecUvrA FKVRDDLTQRLAESFETALELSGGTAVVADMDDP..KAEELLFSANFACPIC.GYSMRELEPRLFSFNNPAGACP
mtUvrA LTVKAAAKRRLTDSVETALNLADGIVVLEFVDHELGAPHREQRFSEKLACPNGHALAVDDLEPRSFSFNSPYGACP
tmUvrA LILETRNEHRILDSLELAMREGKGFVEIRNVD.....TGESKIFSENLMCPVC.GIGFPEITPKLFSFNSPYGACP
slEcm16
180 190 200 210 220 230 240
slEcm16 ACEGIGSVSDIDPAQLF.DASKSLADGAITVPGWKPDGWV.......VQSFTESGFFDPHKAIRDYTEQERHGFLH
snSnorO HCEGLGRVADIDVDQLV.DRERSLNEGAITVPGYAVGSWN.......WQIFAESGLLDPDLRLRDYSPDQWHDLMH
spDrrC DCEGLGHRVQVDLDKLL.DRSKSLNEGAIRHPAFNVGGWF.......WKLYANSGLFDNDKKLRDYTESEWQAFLY
xaUvrA2 TCHGLGRIYDATEATMVLDRSLSIRDRAVAA..WPGA.WH....GQNQRDILTTLGHDVDAPWHTLPKKTRDWILF
drUvrA2 ECHGLGRVYTVTEDSMVPDPSLTIRERAVAA..WPQA.WG....GQNQRDILVTLGIDVDVPWRELPEETRHWILF
ppUvrA2 ECHGMGRVFEVTEATMVPDPSLTIRERAVAA..WPMA.WQ....GQNQRDILVTLGYDVDIPWRDLPQAQRDWILF
bsUvrA DCDGLGVKLEVDLDLVIPNDELTLKEHAIAP..WEPQ.S.SQYYPQLLEAVCRHYGIPMDVPVKELPKEQLAKILY
ecUvrA TCDGLGVQQYFDPDRVIQNPELSLAGGAIRG..WDRR.N..FYYFQMLKSLADHYKFDVEAPWGSLSANVHKVVLY
mtUvrA ECSGLGIRKEVDPELVVPDPDRTLAQGAVAP..WSNG.HTAEYFTRMMAGLGEALGFDVDTPWRKLPAKARKAILE
tmUvrA NCHGLGFTFEVDPSLVIDEEKSVL.EGAIIPYRWDRR.L.....SRWVAREIEKRGVSPHLPFKDLPEDVKEFILY

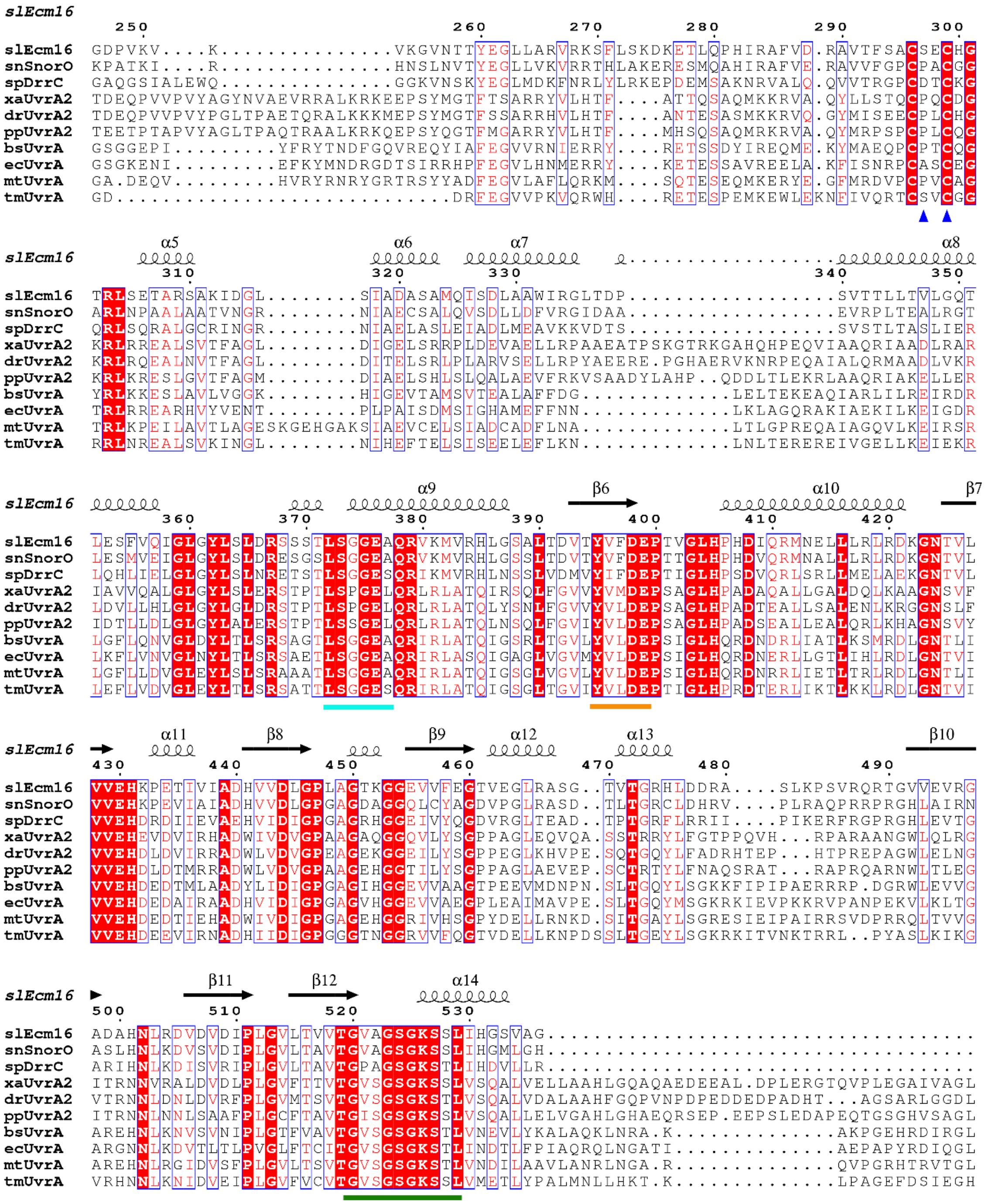
slEcm16
250 260 270 280 290 300
slEcm16 GDPVKV....K...............VKGVNTTYEGLLARVRKSFLSKDKETLQPHIRAFVD.RAVTFSACSECHG
snSnorO KPATKI....R...............HNSLNVTYEGLLVKVRRTHLAKERESMQAHIRAFVE.RAVVFGPCPACGG
spDrrC GAQGSIALEWQ...............GGKVNSKYEGLMDKFNRLYLRKEPDEMSAKNRVALQ.QVVTRGPCDTCKG
xaUvrA2 TDEQPVVPVYAGYNVAEVRRALKRKEEPSYMGTFTSARRYVLHTF....ATTQSAQMKKRVA.QYLLSTQCPQCDG
drUvrA2 TDEQPVVPVYPGLTPAETQRALKKKMEPSYMGTFSSARRHVLHTF....ANTESASMKKRVQ.GYMISEECPLCHG
ppUvrA2 TEETPTAPVYAGLTPAQTRAALKRKQEPSYQGTFMGARRYVLHTF....MHSQSAQMRKRVA.QYMRPSPCPLCQG
bsUvrA GSGGEPI........YFRYTNDFGQVREQYIAFEGVVRNIERRY....RETSSDYIREQME.KYMAEQPCPTCQG
ecUvrA GSGKENI.........EFKYMNDRGDTSIRRHPFEGVLHNMERRY....KETESSAVREELA.KFISNRPCASCEG
mtUvrA GA.DEQV.........HVRYRNRYGRTRSYYADFEGVLAFLQRKM....SQTESEQMKERYE.GFMRDVPCPVCAG
tmUvrA GD...........................DRFEGVVPKVQRWH....RETESPEMKEWLEKNFIVQRTCSVCGG
slEcm16 α5 α6 α7 α8
310 320 330 340 350
slEcm16 TRLSETARSAKIDGL........SIADASAMQISDLAAWIRGLTDP.................SVTTLLTVLGQT
snSnorO ARLNPAALAATVNGR........NIAECSALQVSDLLDFVRGIDAA.................EVRPLTEALRGT
spDrrC QRLSQRALGCRINGR........NIAELASLEIADLMEAVKKVDTS.................SVSTLTASLIER
xaUvrA2 KRLRREALSVTFAGL........DIGELSRRPLDEVAELLRPAAEATPSKGTRKGAHQHPEQVIAAQRIAADLRAR
drUvrA2 KRLRQEALNVTFAGL........DITELSRLPLARVSELLRPYAEERE.PGHAERVKNRPEQAIALQRMAADLVKR
ppUvrA2 KRLKRESLGVTFAGM........DIAELSHLSLQALAEVFRKVSAADYLAHP..QDDLTLEKRLAAQRIAKELLER
bsUvrA YRLKKESLAVLVGGK........HIGEVTAMSVTEALAFFDG.............LELTEKEAQIARLILREIRDR
ecUvrA TRLRREARHVYVENT........PLPAISDMSIGHAMEFFNN.............LKLAGQRAKIAEKILKEIGDR
mtUvrA TRLKPEILAVTLAGESKGEHGAKSIAEVCELSIADCADFLNA.............LTLGPREQAIAGQVLKEIRSR
tmUvrA RRLNREALSVKINGL........NIHEFTELSISEELEFLKN.............LNLTERERREIVGELLKEIEKR
slEcm16 α9 β6 α10 β7
360 370 380 390 400 410 420
slEcm16 LESFVQIGLGYLSLDRSSSTLSGGEAQRVKMVRHLGSALTDVTYVFDEPTVGLHPHDIQRMNELLRLRDKGNTVL
snSnorO LESMVEIGLGYLSLDRESGSLSGGESQRVKMVRHLGSSLTDVTYVFDEPTTGLHPHDVQRMNALLRLRDKGNTVL
spDrrC LQHLIELGLGYLSLNRETSTLSGGESQRIKMVRHLNSSLVDMVYIFDEPTIGLHPSDVQRLSRLLMELAEKGNTVL
xaUvrA2 IAVVQALGLGYLSLERSTPTLSPGELQRLRLATQIRSQLFGVVYVMDEPSAGLHPADAQALLGALDQLKAAGNSVF
drUvrA2 LDVLLHLGLGYLGLDRSTPTLSPGELQRLRLATQLYSNLFGVVYVLDEPSAGLHPADTEALLSALENLKRGGNSLF
ppUvrA2 IDTLLDLGLGYLALERSTPTLSSGELQRLRLATQLNSQLFGVIYVLDEPSAGLHPADSEALLEALQRLKHAGNSVY
bsUvrA LGFLQNVGLDYLTLSRSAGTLSGGEAQRIRLATQIGSRLTGVLYVLDEPSIGLHQRDNDRLIATLKSMRDLGNTLI
ecUvrA LKFLVNVGLNYLTLSRSAETLSGGEAQRIRLASQIGAGLVGVMYVLDEPSIGLHQRDNERLLGTLIHLRDLGNTVI
mtUvrA LGFLLDVGLEYLSLSRAAATLSGGEAQRIRLATQIGSGLVGVLYVLDEPSIGLHQRDNRRLIETLTRLRDLGNTLI
tmUvrA LEFLVDVGLEYLTLSRSATTLSGGESQRIRLATQIGSGLTGVIYVLDEPTIGLHPRDTERLIKTLKKLRDLGNTVI
slEcm16 α11 β8 β9 α12 α13 β10
430 440 450 460 470 480 490
slEcm16 VVEHKPETIVIADHVVDLGPLAGTKGGEVVFEGTVEGLRASG..TVTGRHLDDRA....SLKPSVRQRTGVVEVRG
snSnorO VVEHKPEVIAIADHVVDLGPGAGDAGGQLCYAGDVPGLRASD..TLTGRCLDHRV....PLRAQPRRPRGHLAIRN
spDrrC VVEHDRDIIEVAEHVIDIGPGAGRHGGEIVYQGDVRGLTEAD..TPTGRFLRRII....PIKERFRGPRGHLEVTG
xaUvrA2 VVEHEVDVIRHADWIVDVGPAAGAQGGQVLYSGPPAGLEQVQA.SSTRRYLFGTPPQVH...RPARAANGWLQLRG
drUvrA2 VVEHDLDVIRRADWLVDVGPEAGEKGGEILYSGPPEGLKHVPE.SQTGQYLFADRHTEP...HTPREPAGWLELNG
ppUvrA2 VVEHDLDTMRRADWLVDVGPAAGEHGGTILYSGPPAGLAEVEP.SCTRTYLFNAQSRAT...RAPRQARNWLTLEG
bsUvrA VVEHDEDTMLAADYLIDIGPGAGIHGGEVVAAGTPEEVMDNPN.SLTGQYLSGKKFIPIPAERRRP.DGRWLEVVG
ecUvrA VVEHDEDAIRAADHVIDIGPGAGVHGGEVVAEGPLEAIMAVPE.SLTGQYMSGKRKIEVPKKRVPANPEKVLKLTG
mtUvrA VVEHDEDTIEHADWIVDIGPGAGEHGGRIVHSGPYDELLRNKD.SITGAYLSGRESIEIPAIRRSVDPRRQLTVVG
tmUvrA VVEHDEEVIRNADHIIDIGPGGGTNGGRVVFQGTVDELLKNPDSSLTGEYLSGKRKITVNKTRRL..PYASLKIKG
slEcm16 β11 β12 α14
500 510 520 530
slEcm16 ADAHNLRDVDVDIPLGVLTVVTGVAGSGKSSLIHGSVAG...............................
snSnorO ASLHNLKDVSVDIPLGVLTAVTGVAGSGKSSLIHGMLGH...............................
spDrrC ARIHNLKDISVRIPLGVLTAVTGPAGSGKSTLIHDVLLR...............................
xaUvrA2 ITRNNVRALDVDLPLGVFTTVTGVSGSGKSSLVSQALVELLAAHLGQAQAEDEEAL.DPLERGTQVPLEGAIVAGL
drUvrA2 VTRNNLDNLDVRFPLGVMTSVTGVSGSGKSTLVSQALVDALAAHFGQPVNPDPEDDEDPADHT...AGSARLGGDL
ppUvrA2 ITRNNLNNLSAAFPLGCFTAVTGISGSGKSSLVSQALLELVGAHLGHAEQRSEP.EEPSLEDAPEQTGSGHVSAGL
bsUvrA AREHNLKNVSVNIPLGTFVAVTGVSGSGKSTLVNEVLYKALAQKLNRA.K.............AKPGEHRDIRGL
ecUvrA ARGNNLKDVTLTLPVGLFTCITGVSGSGKSTLINDTLFPIAQRQLNGATI.............AEPAPYRDIQGL
mtUvrA AREHNLRGIDVSFPLGVLTSVTGVSGSGKSTLVNDILAAVLANRLNGA.R.............QVPGRHTRVTGL
tmUvrA VRHNNLKNIDVEIPLGVFVCVTGVSGSGKSSLVMETLYPALMNLLHKT.K.............LPAGEFDSIEGH

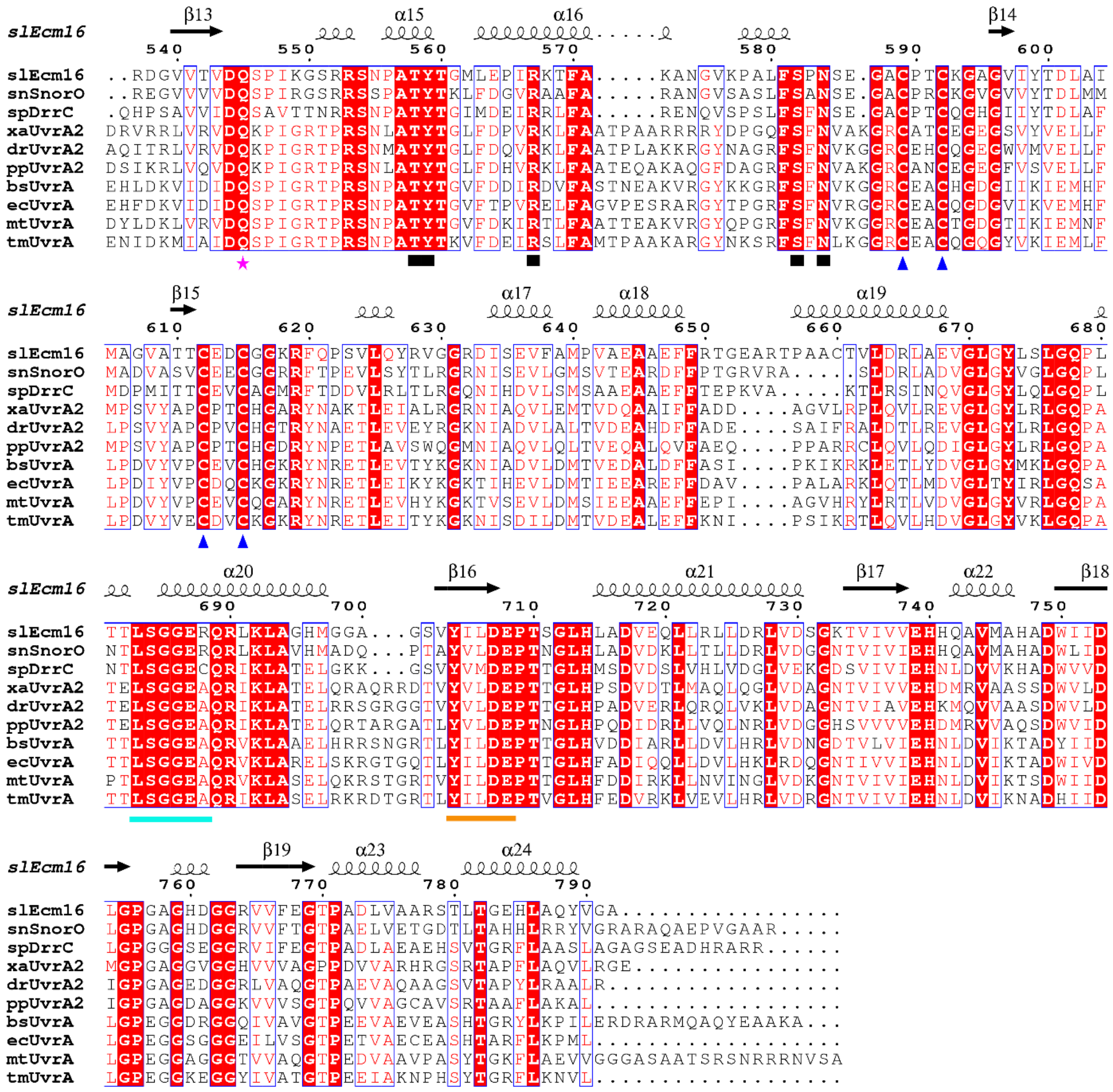


**Figure S1.** Sequence alignment of *S. lasalocidi* Ecm16 (slEcm16), *S. nogalator* SnorO (snSnorO), *S. peuceticus* DrrC (spDrrC), *X. axonopodis* UvrA2 (xaUvrA2), *D. radiodurans* UvrA2 (drUvrA2), *P. putida* UvrA2 (ppUvrA2), *B. stearothermophilus* UvrA (bsUvrA), *E. coli* UvrA (ecUvrA), *M. tuberculosis* UvrA (mtUvrA) and *T. maritima* (tmUvrA). Secondary structure of Ecm16 is noted at the top. Absolutely conserved residues are highlighted in red, and highly conserved residues are enclosed in blue. Conserved DNA-binding residues (black rectangle), Q-loop (magenta star), zinc-coordinating residues (blue triangle), Walker A (green line), Walker B (orange line) and α-helical ABC signature sequence (cyan line) are indicated at the bottom. Sequence alignment was performed using ClustalW (1) and the alignment figure was generated using the ESPript 3.0 program (2).

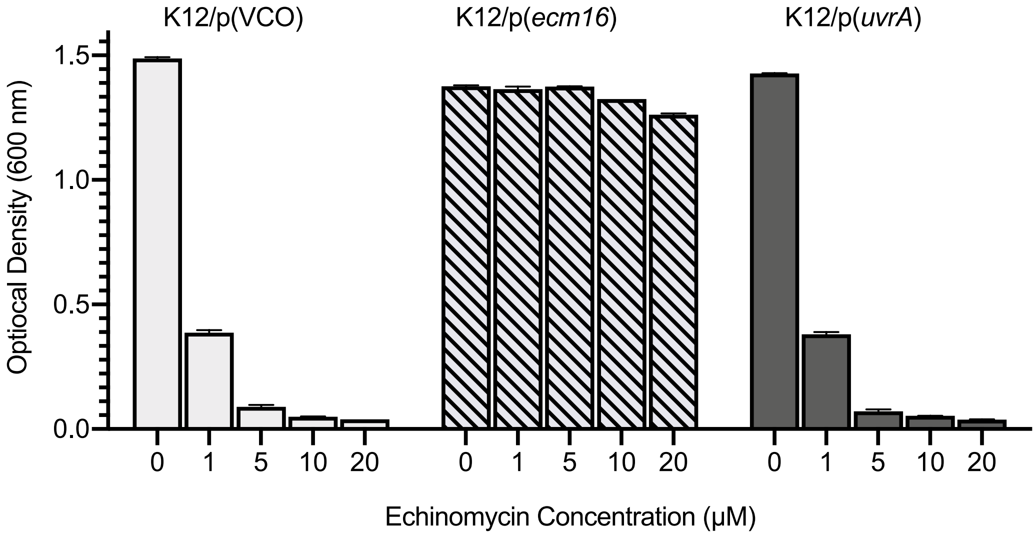


**Figure S2.** Overexpression of *E. coli uvrA* does not confer resistance to echinomycin. Maximum optical densities were determined after 6 h exposure to echinomycin concentrations ranging from 1 μM to 20 μM. Echinomycin was added at time zero to cultures at 0.2 $OD_{600nm}$. *E. coli* K12 strains with pBAD vector-control-only (VCO) or pBAD-*ecm16* (p(*ecm16*) were used for comparison. Error bars represent standard error for duplicate replicates of one trial, analysis is representative of three independent trials.

**Supplementary Tables**

**Table S1. List of strains and plasmids used in this study.**

| Strain # | Description | Genotype | Source |
|---|---|---|---|
| JW4019-2 | K12_Δ*uvrA* | F-, *Δ(araD-araB)567*, *ΔlacZ4787*(::rrnB-3), *λ*, *rph-1*, *Δ(rhaD-rhaB)568*, *ΔuvrA753::kan*, *hsdR514* | (3) |
| JW0762-2 | K12_Δ*uvrB* | F-, *Δ(araD-araB)567*, *ΔlacZ4787*(::rrnB-3), *λ*, *ΔuvrB751::kan*, *rph-1*, *Δ(rhaD-rhaB)568*, *hsdR514* | (3) |
| JW1898-1 | K12_Δ*uvrC* | F-, *Δ(araD-araB)567*, *ΔlacZ4787*(::rrnB-3), *λ*, *ΔuvrC759::kan*, *rph-1*, *Δ(rhaD-rhaB)568*, *hsdR514* | (3) |
| BW25113 | K12 parental strain from Keio collection | F-, *Δ(araD-araB)567*, *ΔlacZ4787*(::rrnB-3), *λ*, *rph-1*, *Δ(rhaD-rhaB)568*, *hsdR514* | (3,4) |
| PM440 | Keio K12 parental strain/pBAD-Myc-HisA | F-, *Δ(araD-araB)567, ΔlacZ4787(::rrnB-3), λ, rph-1, Δ(rhaD-rhaB)568, hsdR514* / pBAD-Myc-HisA | This study |
| PM441 | Keio K12 parental strain/pBAD-Myc-HisA-*ecm16* | F-, *Δ(araD-araB)567*, *ΔlacZ4787*(::rrnB-3), *λ*, *rph-1*, *Δ(rhaD-rhaB)568*, *hsdR514* / pBAD-Myc-HisA-*ecm16* | This study |
| PM442 | K12_Δ*uvrA*/pBAD-Myc-HisA | F-, *Δ(araD-araB)567*, *ΔlacZ4787*(::rrnB-3), *λ*, *rph-1*, *Δ(rhaD-rhaB)568*, *ΔuvrA753::kan*, *hsdR514* /pBAD-Myc-HisA | This study |
| PM443 | K12_Δ*uvrA*/pBAD-Myc-HisA-*ecm16* | F-, *Δ(araD-araB)567*, *ΔlacZ4787*(::rrnB-3), *λ*, *rph-1*, *Δ(rhaD-rhaB)568*, *ΔuvrA753::kan*, *hsdR514* / pBAD-Myc-HisA-*ecm16* | This study |
| PM444 | K12_Δ*uvrB*/pBAD-Myc-HisA | F-, *Δ(araD-araB)567*, *ΔlacZ4787*(::rrnB-3), *λ*, *ΔuvrB751::kan*, *rph-1*, *Δ(rhaD-rhaB)568*, *hsdR514* / pBAD-Myc-HisA | This study |
| PM445 | K12_Δ*uvrB*/pBAD-Myc-HisA-*ecm16* | F-, *Δ(araD-araB)567*, *ΔlacZ4787*(::rrnB-3), *λ*, *ΔuvrB751::kan*, *rph-1*, *Δ(rhaD-rhaB)568*, *hsdR514* / pBAD-Myc-HisA-ecm16 | This study |
| PM446 | K12_Δ*uvrC*/pBAD-Myc-HisA | F-, *Δ(araD-araB)567*, *ΔlacZ4787*(::rrnB-3), *λ*, *ΔuvrC759::kan*, *rph-1*, *Δ(rhaD-rhaB)568*, *hsdR514* / pBAD-Myc-HisA | This study |

| | | | |
|---|---|---|---|
| PM447 | K12_ΔuvrC/pBAD-Myc-HisA-*ecm16* | F-, *Δ(araD-araB)567*, *ΔlacZ4787*(::rrnB-3), *λ*, *ΔuvrC759::kan*, *rph-1*, *Δ(rhaD-rhaB)568*, *hsdR514* */pBAD-Myc-HisA-ecm16* | This study |
| PM483 | K12_Δ*uvrA*/pBAD-*uvrA* | F-, *Δ(araD-araB)567*, *ΔlacZ4787*(::rrnB-3), *λ*, *rph-1*, *Δ(rhaD-rhaB)568*, *ΔuvrA753::kan*, *hsdR514* / pBAD-*uvrA* | This study |
| PM489 | Keio K12 parental strain/pBAD-*uvrA* | F-, *Δ(araD-araB)567*, *ΔlacZ4787*(::rrnB-3), *λ*, *rph-1*, *Δ(rhaD-rhaB)568*, *hsdR514*/ pBAD-*uvrA* | This study |
| PM519 | Keio K12 parental strain/pBAD-*ecm16*-K526A | F-, *Δ(araD-araB)567*, *ΔlacZ4787*(::rrnB-3), *λ*, *rph-1*, *Δ(rhaD-rhaB)568*, *hsdR514/ pBAD-ecm16-K526A* | This study |
| JW3786-5 | K12_Δ*uvrD* | F-, *Δ(araD-araB)567*, *ΔlacZ4787*(::rrnB-3), $\lambda^-$, *rph-1*, *ΔuvrD769::kan*, *Δ(rhaD-rhaB)568*, *hsdR514* | (3) |
| PM535 | K12_Δ*uvrD* / pBAD-Myc-HisA | F-, *Δ(araD-araB)567*, *ΔlacZ4787*(::rrnB-3), $\lambda^-$, *rph-1*, *ΔuvrD769::kan*, *Δ(rhaD-rhaB)568*, *hsdR514/* pBAD-Myc-HisA | This study |
| PM536 | K12_ΔuvrD / pBAD-Myc-HisA-*ecm16* | F-, *Δ(araD-araB)567*, *ΔlacZ4787*(::rrnB-3), $\lambda^-$, *rph-1*, *ΔuvrD769::kan*, *Δ(rhaD-rhaB)568*, *hsdR514/*pBAD-Myc-HisA-*ecm16* | This study |
| **Plasmids** | | | Thermo Fisher |
| pDNA220 | pBAD-Myc-HisA | | Thermo Fisher |
| pDNA223 | pET28a-*ecm16* | | This study |
| pDNA224 | pBAD-Myc-HisA-*uvrA* | | This study |
| pDNA247 | pBAD-Myc-HisA-*ecm16* | | This study |
| pDNA237 | pET28a-*ecm16*-K526A | | This study |
| pDNA239 | pBAD-Myc-HisA-*ecm16*-K526A | | This study |

**Table S2. List of primers used in this study.**

| **Genes** | |
|---|---|
| Ecm16-5'-SacI-fow | AAAGAGCTCCATATGACCGCGGGTACC |
| Ecm16-3'-EcoRI-rev | AAAGAATTCTTACGCGCCCACATACTGCG |
| Ecm16 NdeI-fow | AAACATATGACCGCGGGTACCGAAACCGATACCCAGC |
| Ecm16 EcoRI-rev | AAAGAATTCTTACGCGCCCACATACTGCGCCAGGT |
| Ecm16-BamHI-fow | AAAAATCGCGGATCCATGACCGCGGGTACCGAA A |
| Ecm16-XbaI-rev | AAACGACTAGTCTAGATTACGCGCCCACATACTGCGCC |
| Ecm16K526A-fow | GGTAGCGGTGCAAGCAGCCTGATTCACGGTAGCGTT |
| Ecm16K526A-rev | AATCAGGCTGCTTGCACCGCTACCCGCCA |
| UvrA-SacI-fow | GAGCTCATGGATAAGATCGAAGTTC |
| UvrA-HindIII-rev | AAGCTTTTACAGCATCGGCTTAAG |
| **Sequencing** | |
| T7 promoter | TAATACGACTCACTATAGG |
| T7 terminator | GCTAGTTATTGCTCAGCG |
| pBAD sequencing primer 1-fow (5) | CTGTTTCTCCATACCCGTT |
| pBAD sequencing primer 2 -rev (5) | GGCTGAAAATCTTCTCT |
| pBAD-fow-INVITROGEN | ATGCCATAGCATTTTTATCC |
| pBAD-rev-INVITROGEN | GATTTAATCTGTATCAGG |
| 5'-Mid_ecm16_fow | TAACACCACCTATGAAGGCCTGC |
| 3'-Mid-ecm16-rev | GGTCCACGGTAACCACACCA |
| **Gibson** | |
| Gib_ecm16-down-rev | CTTTTACGAAGAGCGCTCTTCCTTATTACGCGCCCACATACTGC |
| Gib_ecm16-up-fow | TTTGGAGGAGGGCTAGCGAATTCATGACCGCGGGTACCGA |
| Gib_uvrA-pBAD-down-rev | AGTTTTTGTTCGGGCCCAAGCTTTTACAGCATCGGCTTAA |
| Gib_uvrA-pBAD-up-fow | GAATTAACCATGGATCCGAGCTCATGGATAAGATCGAAGT |
| Gib_uvrA-pBAD-up-rev | CCCCGAACTTCGATCTTATCCATGAGCTCGGATCCATGGT |
| Gib_uvrA-pBAD-down-fow | GCTTCCTTAAGCCGATGCTGTAAAAGCTTGGGCCCGAACA |
